# RELEVANCE OF SHAM CONTROL GROUP IN PRECLINICAL ANIMAL STUDIES OF CEREBRAL ISCHEMIA

**DOI:** 10.1101/2023.12.21.572908

**Authors:** María Candamo-Lourido, Esteban López-Arias, Sonia López-Amoedo, Clara Correa-Paz, Susana B. Bravo, Ana Bugallo-Casal, Lucía del Pozo-Filíu, Lara Pérez-Gayol, Nuria Palomar-Alonso, María Pilar Chantada-Vázquez, Francisco Campos, María Pérez-Mato

## Abstract

**Background:** In experimental animal studies, control sham groups are essential to reduce the influence of the surgical intervention on the analysis. The intraluminal filament procedure is one of the most common models of middle cerebral artery occlusion (MCAO) used in the study of cerebral ischemia. However, in these studies, the sham group has not usually been included in the experimental design because of the assumption that the surgical procedure required to access the middle cerebral artery does not affect brain tissue, or that the results obtained from this group are not relevant.

**Objectives:** In this study, we aimed to evaluate the relevance of the sham group by analyzing and comparing the brain protein profile of a sham and an ischemic group subjected to the surgical intraluminal filament occlusion of the middle cerebral artery.

**Material and Methods:** Three randomized experimental groups were tested: control group (healthy animals), sham group, and ischemic group. Twenty-four hours after the interventional procedure, the brain tissue was evaluated by magnetic resonance imaging (MRI). After animal perfusion, the brain is removed for proteomic analysis by liquid chromatography-mass spectrometry (LC-MS/MS) using both a qualitative analysis by data-dependent acquisition (DDA) mode and a quantitative analysis, using a sequential window acquisition of all theoretical mass spectra (SWATH-MS) method on a hybrid quadrupole time-of-flight mass spectrometer.

**Results:** MRI results showed that only animals subjected to cerebral ischemia had ischemic injury. In the sham group 137 dysregulated proteins were detected compared to the 65 in the ischemic group. Moreover, a comparative study of both protein profiles showed the existence of a pool of 17 that appeared dysregulated in both sham and ischemic animals. These results indicate that the surgical procedure required for intraluminal occlusion of the MCA induce changes on brain protein expression that are not associated with the ischemic lesion.

**Conclusion:** This study highlights the importance of including a sham group in the experimental model design to guarantee that the therapeutic target under study is not affected by the surgical intervention.

## Introduction

According to the World Health Organization, stroke is the second most common cause of death and disability in adults worldwide. With the aging of the population, the incidence of stroke in developing countries is expected to continue increasing, becoming the leading cause of premature death and disability in adults. Therefore, the development of safe and effective treatments remains a major challenge for experimental and clinical neuroscience [1].

Animal models are an important tool to study human diseases. In recent decades, several animal models such as the intraluminal monofilament occlusion model, transcranial occlusion, photothrombosis model, thromboembolic occlusion, or endothelin-1-occlusion have been developed for the study of ischemic mechanisms and drug development [2, 3]. However, many of the neuroprotective therapies that showed beneficial effects in preclinical analysis ultimately fail in clinical trials[4]. There are numerous reasons for translational failure in preclinical stroke research conditioned by the experimental setting itself, such as the generation of a plausible hypothesis, the methodological quality of the experimental procedure and the analysis of the results (ideally double-blinded). In order to improve the quality of preclinical studies, the STAIR (Stroke Therapy Academic Industry Roundtable) and ARRIVE (Animal Research: Reporting of in vivo Experiments) guidelines were published [5, 6]. Among the recommendations described, the importance of including a sham group stands out [7]. A sham group can be defined as a study group undergoing a simulated procedure to ensure that they experience the same side effects as the group undergoing the actual surgery or procedure. Sham groups have the potential advantage of reducing the introduction of bias and are used, in experimental designs, to help researchers determine the effectiveness of a drug or treatment and whether a difference between groups is caused by the surgical procedure itself. [8].

Of all the models of cerebral ischemia, the intraluminal model of the middle cerebral artery occlusion (MCAO) is the most widely used, as it allows restoration of blood flow after the induction of focal ischemia, exhibits ischemic penumbra, is highly reproducible, and does not require craniotomy[9]. Despite the characteristics of the surgical procedure of the intraluminal filament model and the recommendation of the ARRIVE guidelines, many studies do not include the sham group in their experimental design. A Pubmed search for “intraluminal filament” in the last 5 years obtained 55 results, while “intraluminal filament and sham group” returned 8, indicating that only 14% of preclinical studies used sham group. In addition to the low percentage of inclusion, in some articles the sham group surgery is not correct, since the procedure consists only on the isolation of the common carotid artery (CCA) from the external carotid artery (ECA), underestimating the effect of the permanent occlusion of the CCA to the brain, required for intraluminal occlusion of the MCA [2, 10]. Therefore, there is no clear consensus on the inclusion of the sham group in experimental designs. To evaluate the impact of surgical procedure on the target organ, we performed an analysis of the brain protein profile of sham and ischemic animals, and the results were compared with a healthy control group without any surgical intervention. Sham control group was submitted to the same surgical procedure but without intraluminal occlusion of the MCA.

With this study, we aim to highlight the importance of using the sham control group of the intraluminal filament model in experimental designs. For this purpose, we performed an analysis of the brain protein profile of sham and ischemic animals compared to control animals by quantitative liquid chromatography–mass spectrometry (LC-MS/MS) by using a sequential window acquisition of all theoretical mass spectra (SWATH-MS) method on a hybrid quadrupole time-of-flight mass spectrometer.

## Results

### MRI analysis of brain tissue in control, sham and ischemic animal

Arterial brain circulation and brain tissue were evaluated in the three groups by MRI. Angiography imaging shows the arterial tree of the brain intact in control animals including the anterior, middle and posteriors cerebral arteries (left and right region), without any signs of ischemic lesion evaluated by T2 imaging at 24h. Sham group was submitted to the same surgical procedure as ischemic animals, without the intraluminal occlusion of the MCA. Angiography imaging indicates that the cerebral vasculature was not altered, and no ischemic lesions were observed either at 24h. In the ischemic group, angiography demonstrated successful occlusion of the MCA that was associated with an ischemic lesion evaluated by T2 imaging at 24h (Fig.1).

**Fig. 1:**
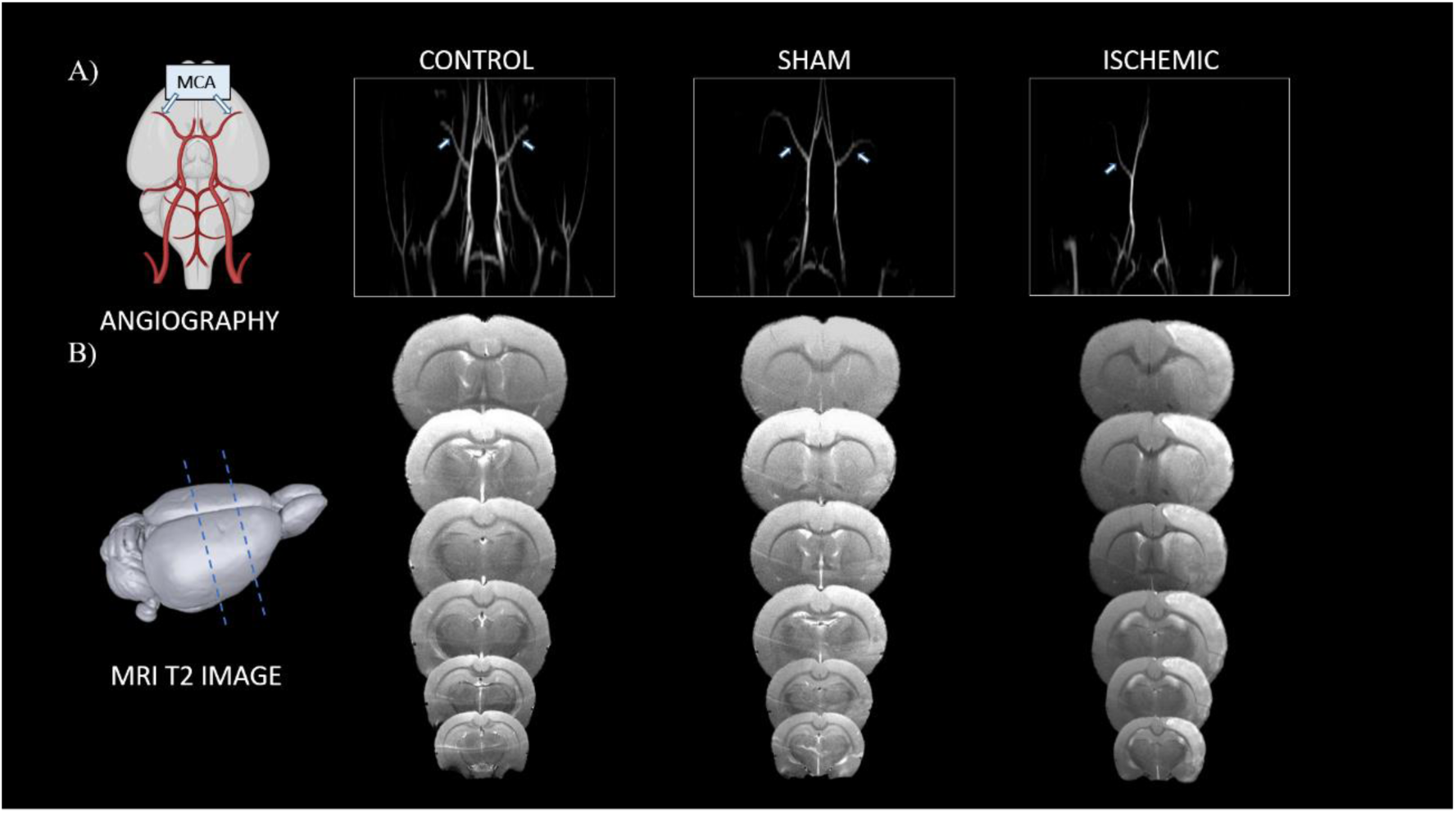
MRI analysis of brain. A) The cerebrovascular anatomy of the rat. Coronal projection of MR angiography image of control, sham and ischemic brain of rats. B) MRI T2 scans of control, sham and ischemic brain of rats at 24 h after tMCAO or surgical procedure. Three animals were included in each group.

### Protein analysis of brain tissue and qualitative protein analysis

Twenty-four hours after surgery, brain tissue of the three experimental groups (control, sham and ischemic group) were analyzed by proteomic analysis. We performed an initial qualitative analysis to identify the set of proteins expressed in each brain tissue using LC-MS/MS technology in data-dependent acquisition (DDA) mode. To obtain representative proteins per group, we selected proteins identified with a false protein discovery rate (FDR)<1% in each sample. Subsequently, quantitative analysis was performed using the SWATH method.

In this study, a total of 2727 proteins were identified with a FDR <1% in each sample. The Venn diagram (Fig.2A) shows the distribution of these proteins and indicates those that were unique and commom to each group. In this analysis, a total of 148 common proteins (Table 1) were detected in the sham and ischemic group, which were subjected to functional analysis using the program Funrich and summarized in Fig.2B according to the biochemical process in which they are involved.

**Fig. 2:**
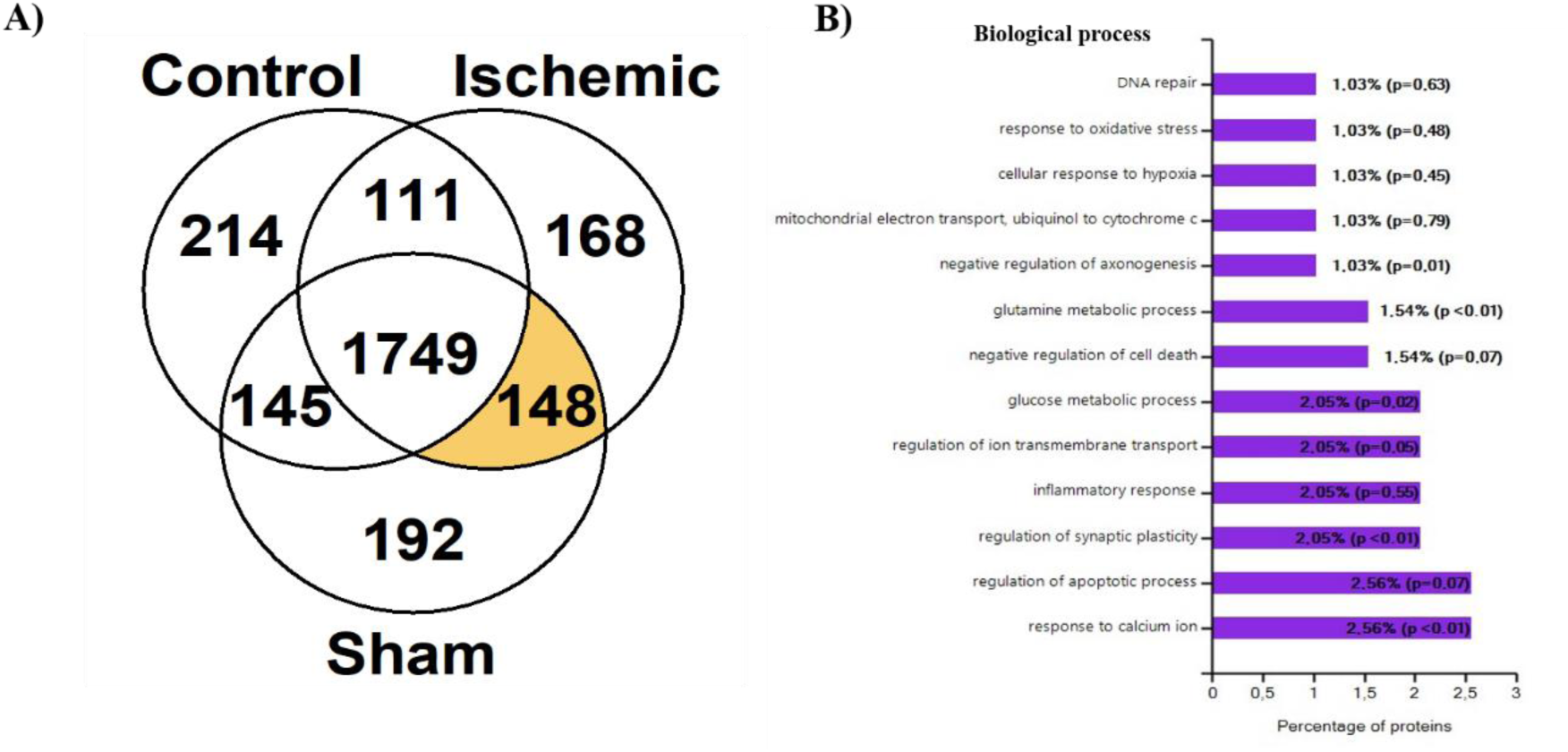
A) Venn diagram showing the distribution of the identified proteins in the three study groups. B) Functional analysis using the FunRich program of common proteins between the sham and ischemic group.

**Table 1:** Common proteins between the sham and ischemic groups identified by qualitative protein analysis. Only proteins with a false discovery rate <1% were selected.

| UnitProt Code | Gene name | Protein Name |
| --- | --- | --- |
| Q6AYZ1 | TBA1C | Tubulin alpha-1C chain |
| P31596-2 | EAA2 | Isoform Glt-1A of Excitatory amino acid transporter 2 |
| Q6URK4-2 | ROA3 | Isoform 2 of Heterogeneous nuclear ribonucleoprotein A3 |
| P14046 | A1I3 | Alpha-1-inhibitor 3 |
| Q03626 | MUG1 | Murinoglobulin-1 |
| Q63041 | A1M | Alpha-1-macroglobulin |
| P09495 | TPM4 | Tropomyosin alpha-4 chain |
| P08010 | GSTM2 | Glutathione S-transferase Mu 2 |
| Q6AXY0 | GSTA6 | Glutathione S-transferase A6 |
| P20059 | HEMO | Hemopexin |
| P29117 | PPIF | Peptidyl-prolyl cis-trans isomerase F, mitochondrial |
| P06399 | FIBA | Fibrinogen alpha chain |
| P24090 | FETUA | Alpha-2-HS-glycoprotein |
| P14480 | FIBB | Fibrinogen beta chain |
| P05545 | SPA3K | Serine protease inhibitor A3K |
| P05544 | SPA3L | Serine protease inhibitor A3L |
| P02767 | TTHY | Transthyretin |
| P01015 | ANGT | Angiotensinogen |
| P04639 | APOA1 | Apolipoprotein A-I |
| P48199 | CRP | C-reactive protein |
| Q9JI12 | VGLU2 | Vesicular glutamate transporter 2 |
| Q63644-2 | ROCK1 | Isoform 2 of Rho-associated protein kinase 1 |
| Q6IE52 | MUG2 | Murinoglobulin-2 |
| E9PTG8 | STK10 | Serine/threonine-protein kinase 10 |
| Q63312 | PHLB1 | Pleckstrin homology-like domain family B member 1 (Fragment) |
| Q9QYP1 | LRP4 | Low-density lipoprotein receptor-related protein 4 |
| P09006 | SPA3N | Serine protease inhibitor A3N |
| D3ZHA0 | FLNC | Filamin-C |
| P15384 | KCNA3 | Potassium voltage-gated channel subfamily A member 3 |
| Q5QD51-2 | AKA12 | Isoform 2 of A-kinase anchor protein 12 |
| P63142 | KCNA2 | Potassium voltage-gated channel subfamily A member 2 |
| P04218 | OX2G | OX-2 membrane glycoprotein |
| P04276 | VTDB | Vitamin D-binding protein |
| P70470 | LYPA1 | Acyl-protein thioesterase 1 |
| Q9QX79 | FETUB | Fetuin-B |
| P19493-2 | GRIA4 | Isoform 2 of Glutamate receptor 4 |
| Q7M733 | HPS6 | Hermansky-Pudlak syndrome 6 protein homolog |
| P14925 | AMD | Peptidylglycine alpha-amidating monooxygenase |
| O08984 | LBR | Delta(14)-sterol reductase LBR |
| P01048 | KNT1 | T-kininogen 1 |
| P10959 | EST1C | Carboxylesterase 1C |
| Q5U2N0 | PYRG2 | CTP synthase 2 |
| Q5M7W4 | TMC5 | Transmembrane channel-like protein 5 |
| D3ZYR1 | FCHO2 | F-BAR domain only protein 2 |
| P63031 | MPC1 | Mitochondrial pyruvate carrier 1 |
| O54874 | MRCKA | Serine/threonine-protein kinase MRCK alpha |
| Q8R508 | FAT3 | Protocadherin Fat 3 |
| P86172 | NMRL1 | NmrA-like family domain-containing protein 1<br>(Fragments) |
| P08934-2 | KNG1 | Isoform LMW of Kininogen-1 |
| Q8CGU9 | TPH2 | Tryptophan 5-hydroxylase 2 |
| O70608 | SYCP2 | Synaptonemal complex protein 2 |
| D4ABL6 | MCTP1 | Multiple C2 and transmembrane domain-containing protein 1 |
| Q9WVR3 | SHIP2 | Phosphatidylinositol 3,4,5-trisphosphate 5-phosphatase 2 |
| P22791 | HMCS2 | Hydroxymethylglutaryl-CoA synthase,<br>mitochondrial |
| Q00566 | MECP2 | Methyl-CpG-binding protein 2 |
| P18292 | THRB | Prothrombin |
| P32198 | CPT1A | Carnitine |
| P20673 | ARLY | Argininosuccinate lyase |
| Q9Z330-4 | DNMT1 | Isoform 4 of DNA (cytosine-5)-methyltransferase<br>1 |
| Q3ZBA0 | TCPR1 | Tectonin beta-propeller repeat-containing protein<br>1 |
| Q09429-2 | ABCC8 | Isoform B of ATP-binding cassette sub-family C<br>member 8 |
| Q5PPG6 | NP1L5 | Nucleosome assembly protein 1-like 5 |
| Q6AXQ4-2 | MTM1 | Isoform 2 of Myotubularin |
| Q4V7C8 | CEP55 | Centrosomal protein of 55 kDa |
| Q5XIT9 | MCCB | Methylcrotonoyl-CoA carboxylase beta chain,<br>mitochondrial |
| Q8R500 | MFN2 | Mitofusin-2 |
| O35430-2 | APBA1 | Isoform 2 of Amyloid-beta A4 precursor protein-<br>binding family A member 1 |
| Q62930 | CO9 | Complement component C9 |
| Q5D006 | UBP11 | Ubiquitin carboxyl-terminal hydrolase 11 |
| P97838-2 | DLGP3 | Isoform 2 of Disks large-associated protein 3 |
| Q53UA7 | TAOK3 | Serine/threonine-protein kinase TA |
| O88277 | FAT2 | Protocadherin Fat 2 |
| Q99PS8 | HRG | Histidine-rich glycoprotein |
| Q9EPY0 | CARD9 | Caspase recruitment domain-containing protein 9 |
| P97519 | HMGCL | Hydroxymethylglutaryl-CoA lyase, mitochondrial |
| P28818 | RGRF1 | Ras-specific guanine nucleotide-releasing factor 1 |
| B2GUZ1 | UBP4 | Ubiquitin carboxyl-terminal hydrolase 4 |
| Q9Z330-2 | DNMT1 | Isoform 2 of DNA (cytosine-5)-methyltransferase 1 |
| Q5PQJ9 | CC175 | Coiled-coil domain-containing protein 175 |
| Q5I0C3 | MCCA | Methylcrotonoyl-CoA carboxylase subunit alpha, mitochondrial |
| P50617 | DEND | Dendrin |
| P69060 | NEUA | N-acylneuraminate cytidyltransferase |
| Q64380 | SARDH | Sarcosine dehydrogenase, mitochondrial |
| P22734 | COMT | Catechol |
| A1L1K8 | HABP4 | Intracellular hyaluronan-binding protein 4 |
| Q5XIV2 | MARHA | Probable E3 ubiquitin-protein ligase MARCHF10 |
| Q63035 | NLRP6 | NACHT, LRR and PYD domains-containing protein 6 |
| Q6T393 | SPT20 | Spermatogenesis-associated protein 20 |
| Q62736 | CALD1 | Non-muscle caldesmon |
| Q6JP77-2 | AKA7G | Isoform Gamma of A-kinase anchor protein 7 isoforms delta and gamma |
| Q5RKH1 | PRP4B | Serine/threonine-protein kinase PRP4 homolog |
| Q02293 | FNTB | Protein farnesyltransferase subunit beta |
| D3ZV31 | ZCHC4 | rRNA N6-adenosine-methyltransferase ZCCHC4 |
| P49303-2 | POLG | Isoform Lb of Genome polyprotein |
| P55266 | DSRAD | Double-stranded RNA-specific adenosine deaminase |
| O88943-3 | KCNQ2 | Isoform C of Potassium voltage-gated channel subfamily KQT member 2 |
| D3Z8N4 | KLH20 | Kelch-like protein 20 |
| Q69CM7-2 | MYBPP | Isoform 2 of MYCBP-associated protein |
| Q5U2R7 | MESD | LRP chaperone MESD |
| Q5M7W7 | SYPM | Probable proline--tRNA ligase, mitochondrial |
| P24392 | PEX2 | Peroxisome biogenesis factor 2 |
| P97610 | SYT12 | Synaptotagmin-12 |
| Q6MGA9 | BRD2 | Bromodomain-containing protein 2 |
| Q5PPN7 | MITOK | Mitochondrial potassium channel |
| Q8CF82 | ABCA5 | Cholesterol transporter ABCA5 |
| Q80ZF0 | CORA1 | Collagen alpha-1(XXVII) chain |
| Q9JHZ9 | S38A3 | Sodium-coupled neutral amino acid transporter 3 |
| P16067 | ANPRB | Atrial natriuretic peptide receptor 2 |
| Q157S1 | CRTC1 | CREB-regulated transcription coactivator 1 |
| Q99068 | AMRP | Alpha-2-macroglobulin receptor-associated protein |
| Q02765 | CATS | Cathepsin S |
| P22199-3 | MCR | Isoform 3 of Mineralocorticoid receptor |
| Q6AYT7 | ABD12 | Lysophosphatidylserine lipase ABHD12 |
| Q5XIU1 | ABTB1 | Ankyrin repeat and BTB/P |
| P01836 | KACA | Ig kappa chain C region, A allele |
| O70467 | ANM3 | Protein arginine N-methyltransferase 3 |
| B4F785 | EGFLA | Pikachurin |
| Q9WTW1-2 | GRIP2 | Isoform 2 of Glutamate receptor-interacting protein 2 |
| Q4KM33 | PLEK | Pleckstrin |
| Q68FT3 | PYRD2 | Pyridine nucleotide-disulfide oxidoreductase domain-containing protein 2 |
| Q810H6-2 | DISC1 | Isoform 2 of Disrupted in schizophrenia 1 homolog |
| Q6QD51 | CCD80 | Coiled-coil domain-containing protein 80 |
| Q6MG55 | ABHGA | Phosphatidylserine lipase ABHD16A |
| Q4KM84 | MET18 | Histidine protein methyltransferase 1 homolog |
| Q9QWS8 | KCNH8 | Potassium voltage-gated channel subfamily H member 8 |
| Q3LUD4 | LZTS2 | Leucine zipper putative tumor suppressor 2 |
| P50609 | FMOD | Fibromodulin |
| P14270-6 | PDE4D | Isoform 9 of cAMP-specific 3',5'-cyclic phosphodiesterase 4D |
| Q9QZU7 | BODG | Gamma-butyrobetaine dioxygenase |
| P09367-2 | SDHL | Isoform 2 of L-serine dehydratase/L-threonine deaminase |
| Q4V7C4 | CCD71 | Coiled-coil domain-containing protein 71 |
| P97527 | CNTN5 | Contactin-5 |
| Q9JMI1 | AACS | Acetoacetyl-CoA synthetase |
| Q99M76 | STAG3 | Cohesin subunit SA-3 |
| P62275 | RS29 | 40S ribosomal protein S29 |
| Q6XQN1 | PNCB | Nicotinate phosphoribosyltransferase |
| Q9ERE4 | GOLP3 | Golgi phosphoprotein 3 |
| Q9WVJ4 | SYJ2B | Synaptojanin-2-binding protein |
| Q62947-2 | ZEB1 | Isoform 2 of Zinc finger E-box-binding homeobox 1 |
| P43165 | CAH5A | Carbonic anhydrase 5A, mitochondrial |
| D3ZGQ5 | NEK8 | Serine/threonine-protein kinase Nek8 |
| Q5R JL0-2 | ERMIN | Isoform 2 of Ermin |
| D3ZBM7 | HACE1 | E3 ubiquitin-protein ligase HACE1 |
| Q6RFY2 | PHAR3 | Phosphatase and actin regulator 3 |
| Q62789 | UD2B7 | UDP-glucuronosyltransferase 2B7 |
| Q6AYQ4 | TM109 | Transmembrane protein 109 |
| P62850-2 | RS24 | Isoform 2 of 40S ribosomal protein S24 |
| Q5XIE1 | THEM6 | Protein THEM6 |

### Quantitative protein analysis by SWATH

Quantitative analysis was subsequently performed using the SWATH method to compare the protein profile expression of sham vs control and ischemia vs control. Dysregulated proteins were identified when p < 0.05, and fold change (FC) > 1.5 or <0.8 were chosen as cut-offs.

In the first analysis (sham vs control), a total of 137 dysregulated proteins were identified, of which 55 were downregulated and 87 upregulated in the sham group with respect to the control (Table 2). Using the String software to determine the biological processes associated with those dysregulated proteins we found that they are involved in calcium-ion regulated exocytosis, synaptic vesicle cycle, neurotransmitter secretion, glial cell differentiation, cellular response to oxidative stress, regulation of neuron death and apoptotic process, response to stress, and regulation of metabolic process (Fig. 3A). The volcano plot (Fig.3B) shows the differences in protein expression between the sham-control group. Among the major dysregulated proteins are ECI2, PI42C, HNRPL, HPCL4, TBA4A, NFH, S6A11, GPM6B, CSN1, HBB2, HBB1 and HBA, which were upregulated more than 700-fold; the downregulated were reduced in this comparison group by 26 fold (Fig. 3C, D).

**Fig. 3:**
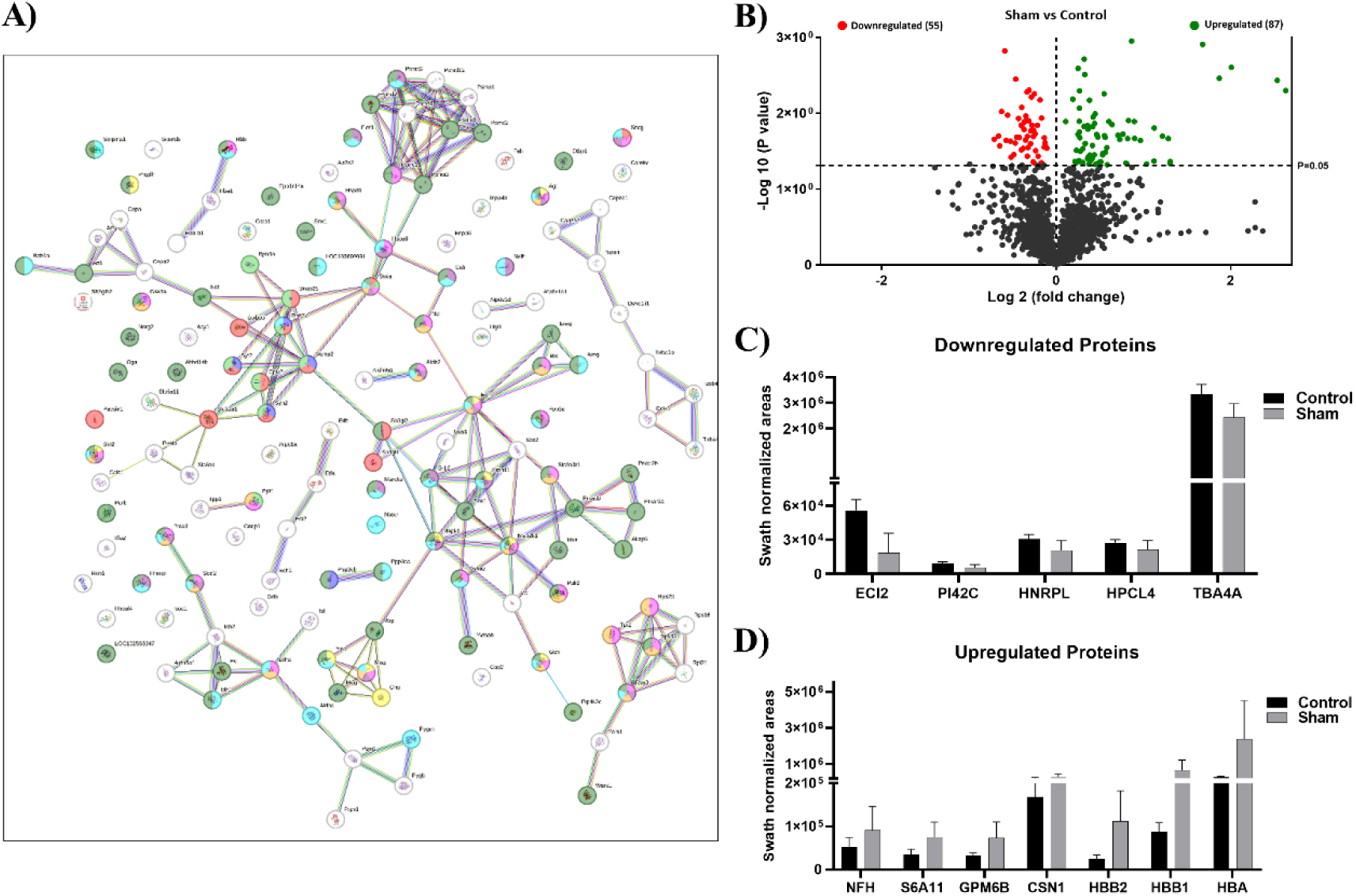
Dysregulated proteins Sham vs. Control. A) STRING interaction analysis of dysregulated proteins. Red dots indicate proteins related to regulation of neuron death and apoptotic process; green dots represent proteins involved in stress and regulation of metabolic process; and blue dots associate with cellular response to oxidative stress. B) Volcano plot resulting from comparison of sham and control. The cut-off for protein expression changes was p < 0.05, and fold change (FC) > 1.5 or <0.8. Up- and downregulated dysregulated proteins are represented as green and red dots, respectively. Proteins that were not differentially expressed are presented as black dots. C, D) Top 12 proteins that exhibited the greatest fold change in expression in Sham vs. Control. The data are presented as the mean ± standard deviation (SD) of the mean. Three animals were included in each comparison group.

**Table 2:**
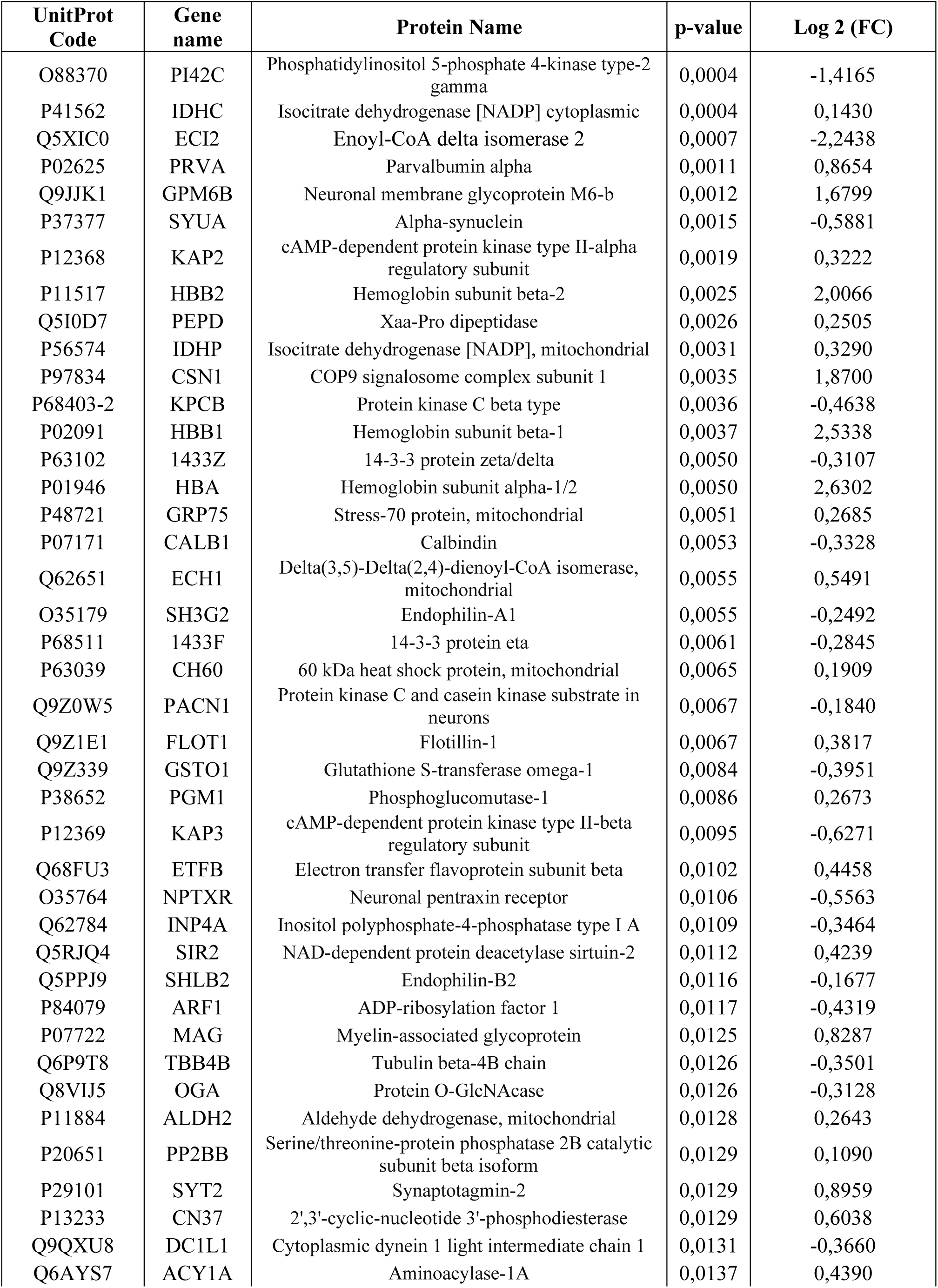
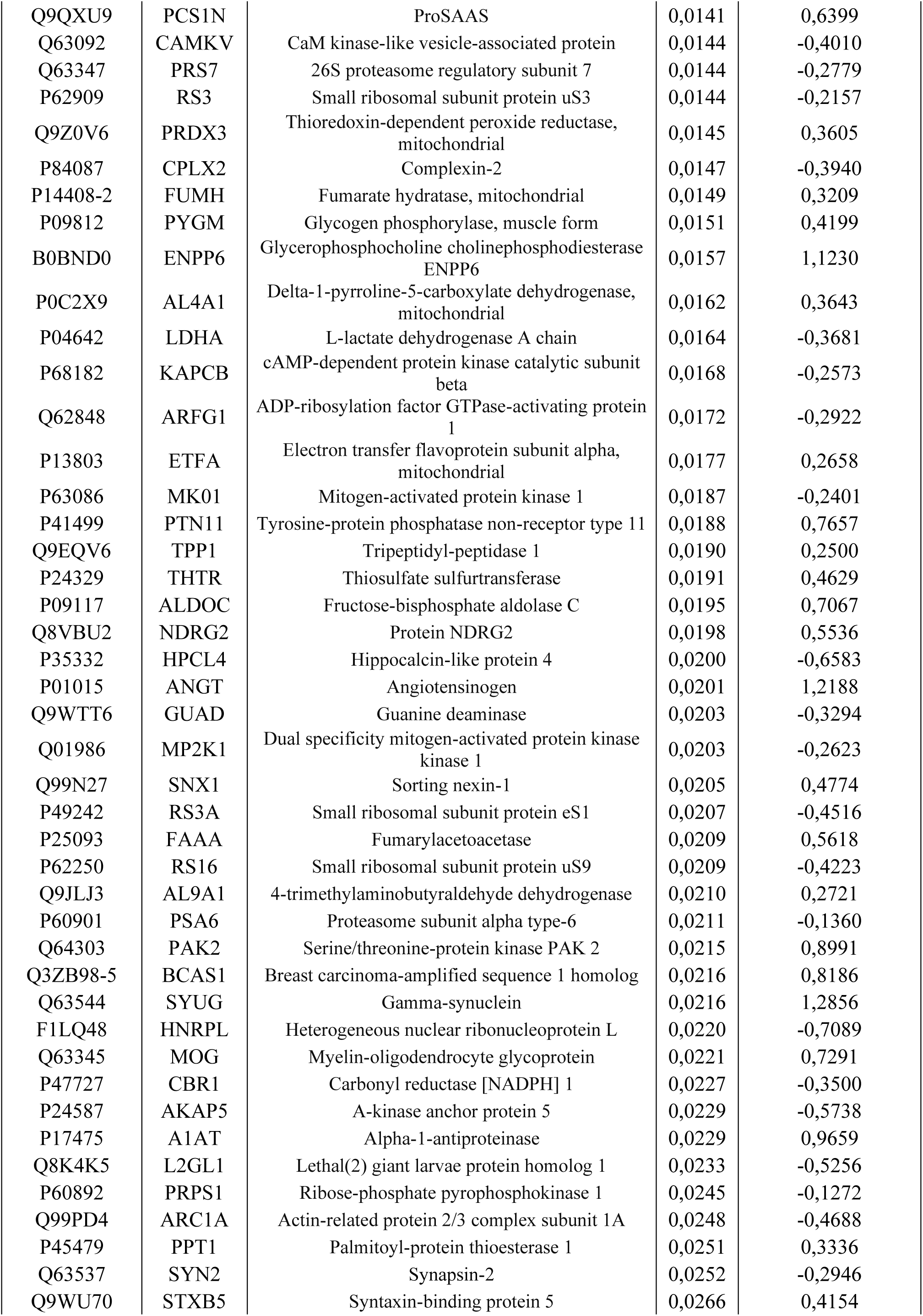
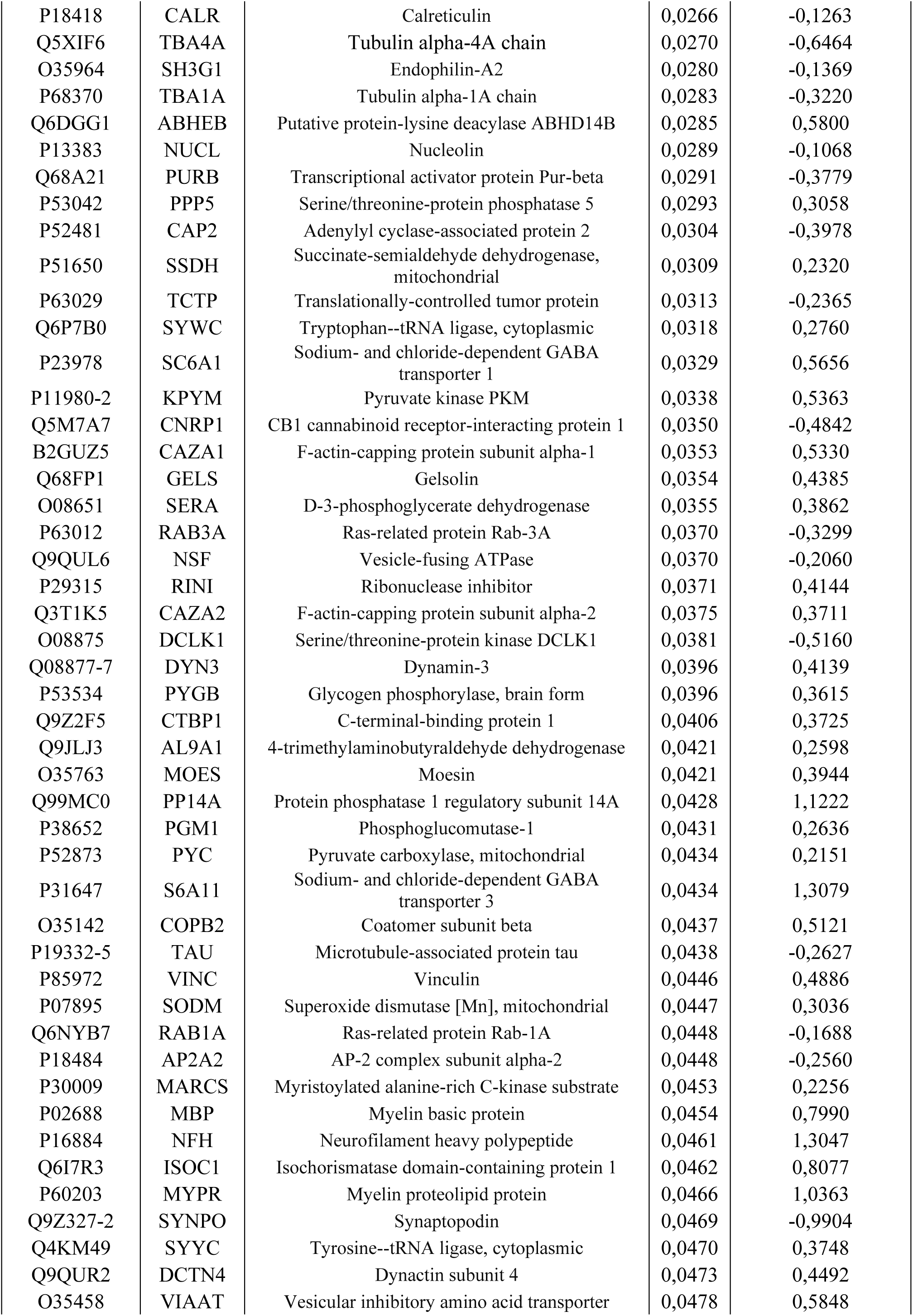
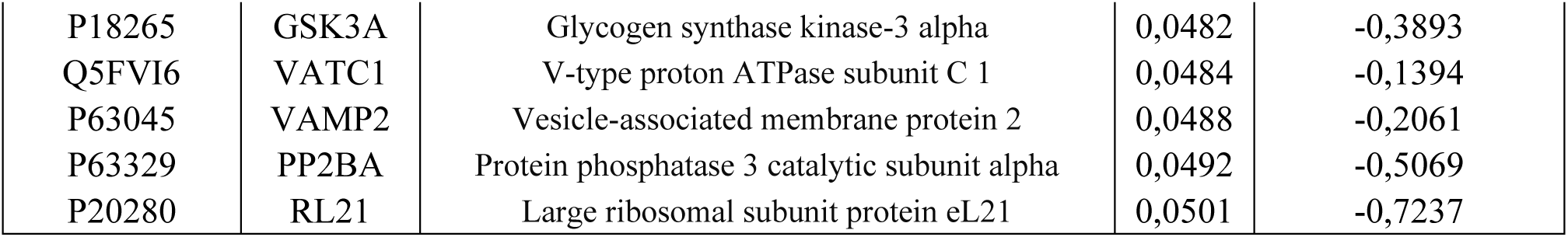
Dysregulated proteins Sham vs. Control. Proteins considered dysregulated are those with a p-value < 0.05 and a Log 2 Fold Change (FC) > 1.5 or <0.8.

Comparative protein analysis between the ischemic and control groups determined a total of 65 dysregulated proteins, 35 downregulated and 30 upregulated (Table 3). The String Pathway showed that dysregulated proteins were implicated in the following biological processes: regulation of telomere maintenance via telomerase, negative regulation of ubiquitin-dependent protein catabolic process, regulation of DNA biosynthetic process, protein stabilization, regulation of proteolysis and catabolic process and cellular metabolic process (Fig. 4A). The differences in protein expression between ischemia vs control are shown in the volcano diagram (Fig. 4B). The most important upregulated proteins were ACADL, HBB1 and HBA, which exhibited an increase of 400-fold in their expression in the ischemic tissue. At the same time, SYUG, PP1A, FRIL1, RTN1, ARP5L, MGLL, PCP4, NCKP1 showed a decrease in 40-fold in the ischemic group with respect to the control (Fig. 4C, D).

**Fig. 4:**
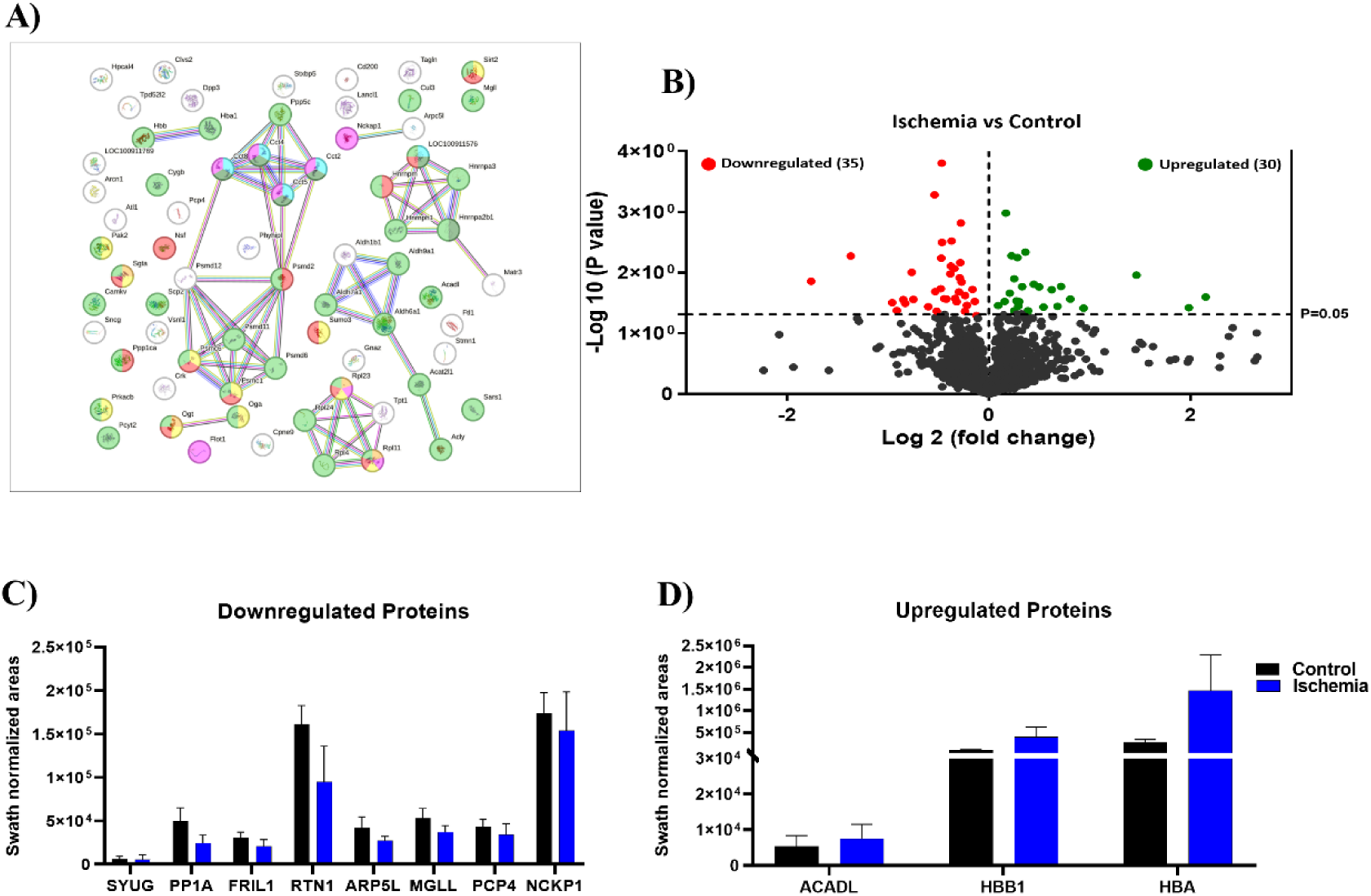
Dysregulated proteins Ischemia vs. Control. A) STRING interaction analysis of dysregulated proteins. Pink dots indicate proteins related with catabolic process; green dots represent proteins involved in regulation of DNA biosynthetic process; and yellow dots are associated with regulation of proteolysis and catabolic process and cellular metabolic process. B) Volcano plot resulting from comparison of ischemia and control. The cut-off for protein expression changes was p < 0.05, and fold change (FC) > 1.5 or <0.8. Up- and downregulated dysregulated proteins are represented as green and red dots, respectively. Proteins that were not differentially expressed are presented as black dots. C, D) Top 11 proteins that exhibited the greatest fold change in expression in Ischemia vs. Control. The data are presented as the mean ± standard deviation (SD) of the mean. Three animals were included in each comparison group.

**Table 3:**
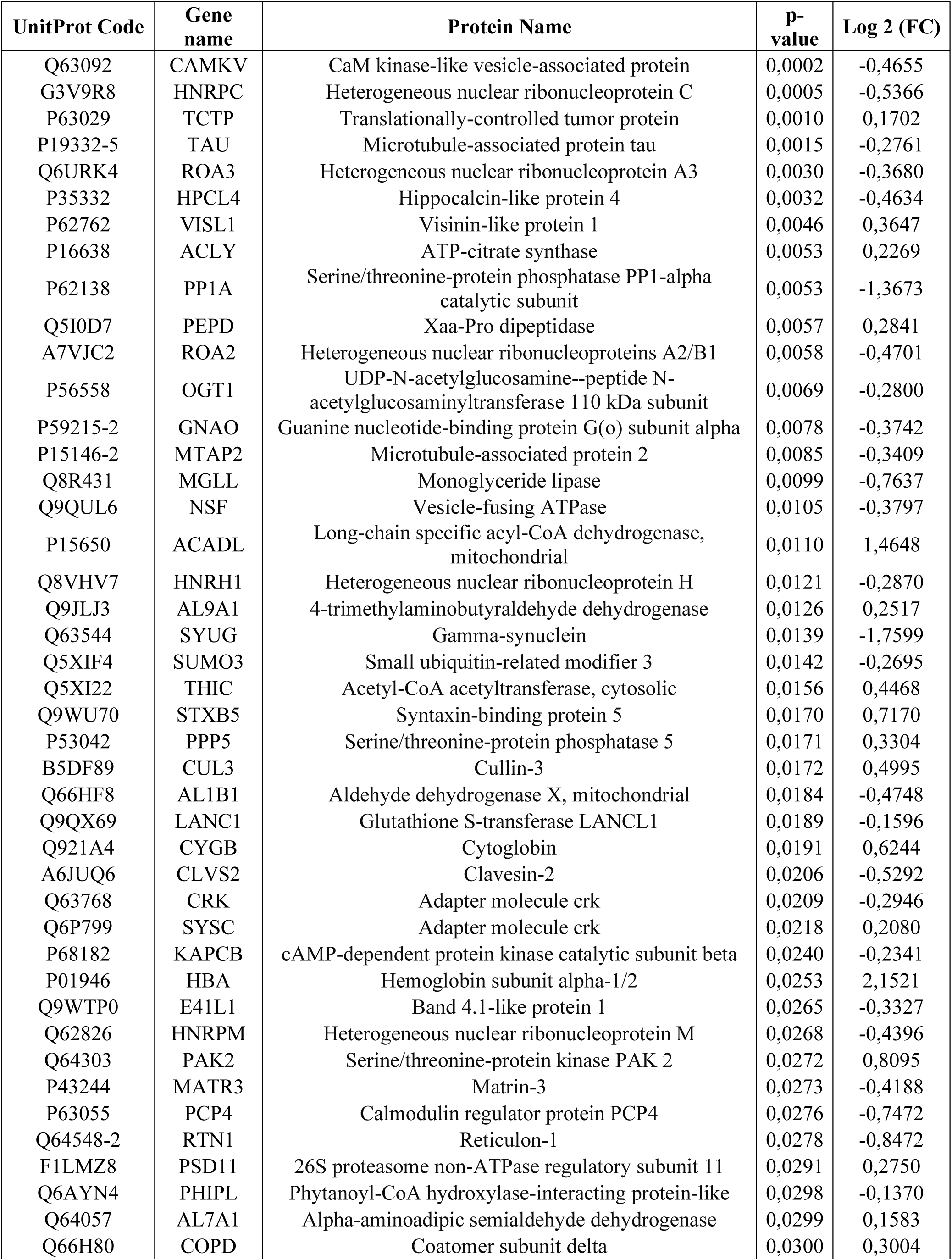
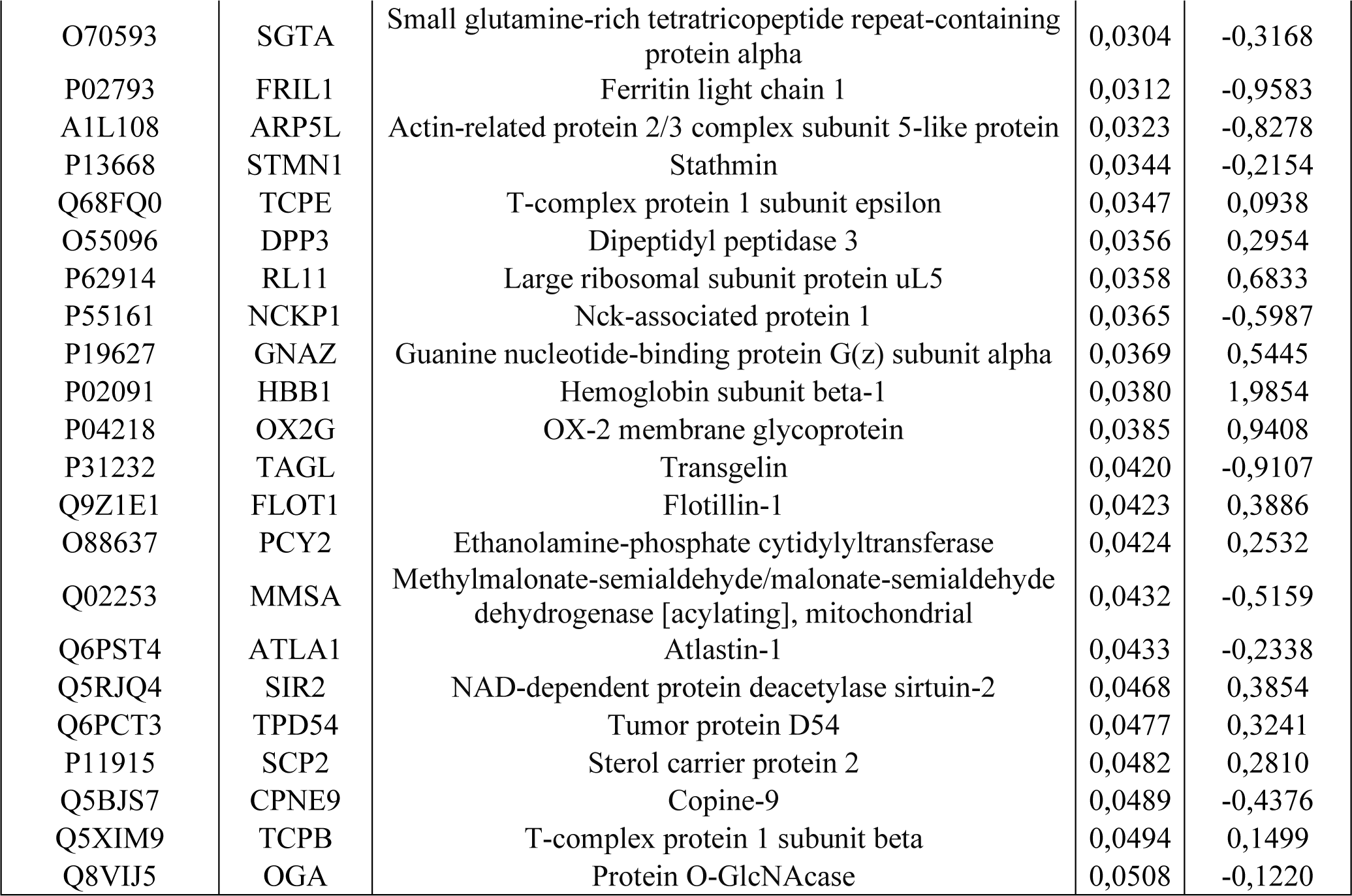
Dysregulated proteins Ischemic vs. Control. Proteins considered dysregulated are those with a p-value < 0.05 and a Log 2 Fold Change (FC) > 1.5 or <0.8.

Finally, we performed an analysis of the common dysregulated proteins in the sham and ischemic groups. Venn diagram (Fig. 5A) shows the distribution of proteins in each of the groups, indicating that there are 17 common proteins that appeared affected in both groups (Table 4, 5). With the exception of SYUG and TCTP proteins, all other proteins are dysregulated in the same way in the sham and ischemic groups. Thus, PEPD, AL9A1, PPP5, FLOT1, STXB5, SIR2, PAK2, HBB1, HBA are upregulated, whereas HPCL4, CAMKV, OGA, TAU, KAPCB, NSF are downregulated in both groups (Fig. 5B, C). A functional analysis of these proteins was performed using the Funrich and Reactome programs. The upregulated proteins are mainly involved in the following biological processes: amino acid metabolism (PEPD, AL9A1), apoptotic processes and cell death (PPP5, FLOT1, PAK2), calcium-dependent exocytosis and neurotransmitter release (STXB5), autophagy (SIR2), and oxygen transport from the lungs to the various peripheral tissues (HBB1, HBA). As far as downregulated proteins are concerned, they play a fundamental role in: calcium-dependent pathway regulation (HPCL4), association with the plasma membrane of soma and in neurites, axons and dendrites (CAMKV), ischemia-reperfusion-related apoptosis and cell death (OGA), maintenance of neuronal polarity (TAU), synaptic plasticity (KAPCB), and catalyzation of the fusion of transport vesicles within the Golgi cisternae (NSF). SYUG and TCTP proteins are involved in neurofilament network integrity and calcium binding and microtubule stabilization respectively.

**Fig. 5:**
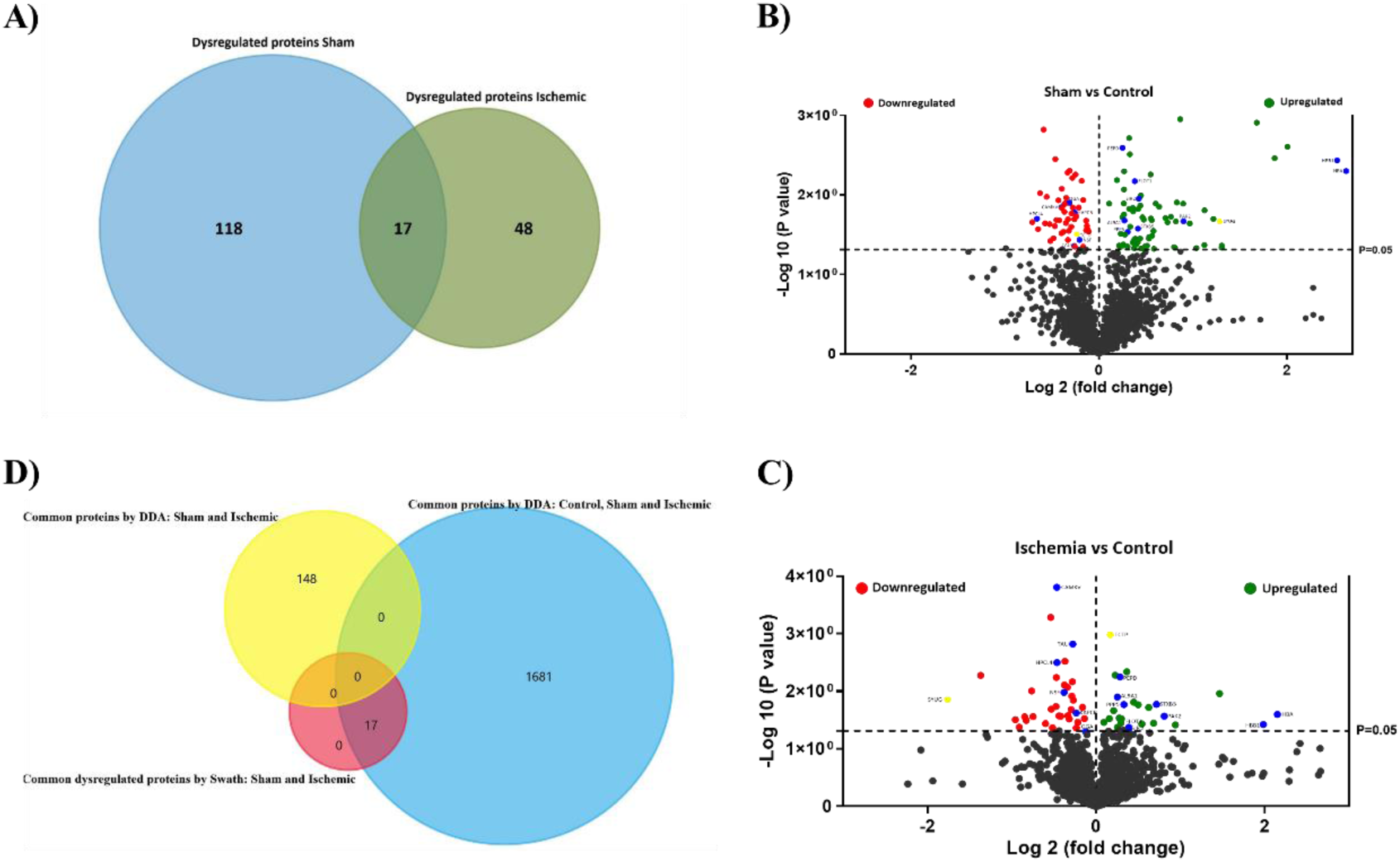
Common dysregulated proteins between sham and ischemic group. A) Venn diagram showing the overlap of common dysregulated proteins between sham and ischemic groups. B, B) Results of the dysregulated proteins presented correspond to Sham vs. control (B) and Ischemia vs. control (C) using volcano plots. The cut-off point for protein expression changes was p < 0.05, and fold change (FC) > 1.5 or <0.8. Up- and down-regulated proteins are represented as green and red dots, respectively. Common dysregulated proteins in the sham and ischemic groups are represented as blue dots. Yellow dots represent SYUG and TCTP proteins, which are the common proteins dysregulated differently. Proteins that were not differentially expressed are presented as black dots. D) Venn diagram showing the overlap of common dysregulated proteins identified by swath, common proteins between sham and ischemia by DDA, and the protein pool common to control, sham and ischemic detected by DDA. Three animals were included in each comparison group.

**Table 4:** Common dysregulated proteins in the sham group. Proteins considered dysregulated are those with a p-value < 0.05 and a Fold Change (FC) > 1.5 or <0.8.

| UniProt Code | Gene name | Protein Name | P-value | Fold Change |
| --- | --- | --- | --- | --- |
| Q5I0D7 | PEPD | Xaa-Pro dipeptidase | 0,0026 | 1,1896 |
| P02091 | HBB1 | Hemoglobin subunit beta-1 | 0,0037 | 5,7908 |
| P01946 | HBA | Hemoglobin subunit alpha-1 | 0,0050 | 6,1912 |
| Q9Z1E1 | FLOT1 | Flotillin-1 | 0,0067 | 1,3029 |
| Q5RJQ4 | SIR2 | NAD-dependent protein deacetylase sirtuin-2 | 0,0112 | 1,3415 |
| Q8VIJ5 | OGA | Protein O-GlcNAcase | 0,0126 | 0,8051 |
| Q63092 | CAMKV | CaM kinase-like vesicle-associated protein | 0,0144 | 0,7574 |
| P27791 | KAPCB | cAMP-dependent protein kinase catalytic subunit alpha | 0,0168 | 0,8366 |
| P35332 | HPCL4 | Hippocalcin-like protein 4 | 0,0200 | 0,6336 |
| Q64303 | PAK2 | Serine threonine-protein kinase PAK 2 | 0,0215 | 1,8649 |
| Q63544 | SYUG | Gamma-synuclein | 0,0216 | 2,4379 |
| Q9WU70 | STXB5 | Syntaxin-binding protein 5 | 0,0266 | 1,3337 |
| P53042 | PPP5 | Serine threonine-protein phosphatase 5 | 0,0293 | 1,2361 |
| P63029 | TCTP | Translationally-controlled tumor protein | 0,0313 | 0,8488 |
| Q9QUL6 | NSF | Vesicle-fusing ATPase | 0,0370 | 0,8670 |
| Q9JLJ3 | AL9A1 | 4-trimethylaminobutyraldehyde dehydrogenase | 0,0421 | 1,1973 |
| P19332-5 | TAU | Isoform Tau-E of Microtubule-associated protein tau | 0,0438 | 0,8335 |

**Table 5:** Common dysregulated proteins in the ischemic group. Proteins considered dysregulated are those with a p-value < 0.05 and a Fold Change (FC) > 1.5 or <0.8.

| UniProt Code | Gene name | Protein Name | P-value | Fold Change |
| --- | --- | --- | --- | --- |
| Q63092 | CAMKV | CaM kinase-like vesicle-associated protein | 0,0002 | 0,7242 |
| P63029 | TCTP | Translationally-controlled tumor protein | 0,0010 | 1,1252 |
| P19332-5 | TAU | Isoform Tau-E of Microtubule-associated protein tau | 0,0015 | 0,8258 |
| P35332 | HPCL4 | Hippocalcin-like protein 4 | 0,0032 | 0,7253 |
| Q5I0D7 | PEPD | Xaa-Pro dipeptidase | 0,0057 | 1,2177 |
| Q9QUL6 | NSF | Vesicle-fusing ATPase | 0,0105 | 0,7686 |
| Q9JLJ3 | AL9A1 | 4-trimethylaminobutyraldehyde dehydrogenase | 0,0126 | 1,1907 |
| Q63544 | SYUG | Gamma-synuclein | 0,0139 | 0,2953 |
| Q9WU70 | STXB5 | Syntaxin-binding protein 5 | 0,0170 | 1,6438 |
| P53042 | PPP5 | Serine threonine-protein phosphatase 5 | 0,0171 | 1,2574 |
| P27791 | KAPCB | cAMP-dependent protein kinase catalytic subunit alpha | 0,0240 | 0,8502 |
| P01946 | HBA | Hemoglobin subunit alpha-1 | 0,0253 | 4,4448 |
| Q64303 | PAK2 | Serine threonine-protein kinase PAK 2 | 0,0272 | 1,7526 |
| P02091 | HBB1 | Hemoglobin subunit beta-1 | 0,0380 | 3,9596 |
| Q9Z1E1 | FLOT1 | Flotillin-1 | 0,0423 | 1,3091 |
| Q5RJQ4 | SIR2 | NAD-dependent protein deacetylase sirtuin-2 | 0,0468 | 1,3062 |
| Q8VIJ5 | OGA | Protein O-GlcNAcase | 0,0508 | 0,9189 |

Besides, we analyzed whether the dysregulated common proteins were detected by qualitative proteomic analysis. For this purpose, we performed a comparison between the dysregulated common proteins in the sham and ischemic groups identified by SWATH, with the pool of common proteins in the control, sham and ischemic groups (1749 proteins see Fig. 2A), and the common proteins between the sham and ischemic groups (148 proteins see Fig. 2A) identified by the qualitative method. These 17 proteins common to the sham and ischemic groups were also found within the pool of proteins common to all three groups (Fig. 5D).

## DISCUSSION

The occlusion of the MCA is one of the most common vessels affected in clinical ischemic stroke, and therefore, the intraluminal occlusion of this major vessel is the most used experimental approach in animal models to replicate the clinical condition. The intraluminal filament occlusion of the MCA animal model was initially developed by Koizumi in 1986 and modified by Longa in 1989 [11]. In this model, the access to the MCA requires to introduce the filament into the internal carotid artery through the external carotid. Infarcts induced by this approach often comprise both striatal and cortical damage. Temporary insertion of a filament in the MCA, which is later removed after the desired period of ischemia, produces a transient MCA territory ischemia followed by the restoration of blood circulation. Alternatively, leaving the filament in the MCA can be used to reproduce a permanent stroke without reperfusion.

The main advantages of this method include the reliability to reproduce the pathophysiology of stroke, the ability to avoid craniotomy and the potential to develop high-throughput drug screenings. However, although this is the most common model used in experimental stroke field, the main disadvantages include tracheal edema, paralysis of muscles of mastication and swallowing caused by the injury to access to the CCA, ICA and ECA. In addition, the clamping of the carotid arteries, necessary to introduce the filament into the vessels, causes a reduction in cerebral flow, which could contribute to brain injury beyond the ischemia caused by the occlusion of the MCA. Considering that the surgical procedure itself can interfere with the final results, it is particularly relevant to include the sham control group [12]. There are studies that support this premise, as they have observed that surgical procedures are not harmless to the tissue. Thus, in a study on liver regeneration [12], the effects of sham surgery procedures and anesthesia was analyzed on the expression pattern of microRNAs in rat liver. They found 49 microRNAs modified by hepatectomy and 45 modified by sham laparotomy, with 10 microRNAs showing similar changes after both real and sham surgery. The impact of sham surgery has also been highlighted by Cole et al.[13], who compared the effects of standard sham procedures used in research on traumatic brain injury (craniotomy by drill or manual trepanation) with the effects of anesthesia alone. They found that the traditional sham control induced significant pro-inflammatory, morphological and behavioral changes, and that these could confound the interpretation in brain injury models.

In line with these previous facts, qualitative analysis evidenced the existence of 192 proteins that are exclusively modified in the sham group and other 148 common proteins that are altered in both sham and ischemic groups. The quantitative analysis sham vs control showed a total of 137 deregulated proteins, while ischemia vs control 65 deregulated proteins. These data are in accordance with Rutkai et al [14] studies, who describe that variations in cerebrovascular function and mitochondrial bioenergetics occur in the sham group of MCAO. However, most studies that performed proteomics of tissue from the MCAO sham group found no proteome modifications [15, 16]. This may be because those works assume that sham is a fully healthy control when a non-interventional control group should be included in the analysis, as we have done in this study. In our analysis it was also observed that the number of dysregulated proteins was higher in the sham group than in the ischemic group, which could be related to the degradation of proteins in the ischemic region.

We later performed an analysis of common dysregulated proteins in the sham and ischemic group and the following common proteins were found: SYUG, TCTP, PEPD, AL9A1, PPP5, FLOT1, STXB5, SIR2, PAK2, HBB1, HBA, HPCL4, CAMKV, OGA, TAU, KAPCB and NSF. Most of them have been described to be involved in cerebral ischemia: SYUG was detected in patients with chronic cerebral ischemia [14]; PEPD is involved in the glutamatergic regulatory system of cerebral ischemia [15]; AL9A1 was found in the neuroproteome of ischemic animals [16]; PPP5 is involved in the ischemia-induced tyrosine phosphorylation of proteins [17]; FLOT1 protects cortical neurons against hypoxia/reoxygenation injury [11]; STXB5 participates in the blood-brain barrier breakdown and activation of microglia [18]; SIR2 was upregulated during neuronal ischemia and translocated into neuronal nuclei, while downregulation of SIR2 could significantly protect neurons against cerebral ischemia [19]; HBB1 and HBA were up-regulated during hypoxia [20]; CAMKV participates in processes of repair or replenishment of neuronal circuits during recovery from brain damage [21]; OGA ameliorate cerebral ischemia-reperfusion injury [22]; and TAU contributes to progressive damage in cerebral ischemia, likely by regulating excitotoxic signaling [23].

Beyond the molecular mechanism in which these dysregulated proteins could be involved in, these common affected proteins in both sham and ischemic indicates that surgical intervention of the MCAO model induces changes on brain proteins that could lead to an overestimation of the molecular mechanisms caused by the ischemic damage.

In conclusion, it is widely assumed that surgical intervention to access the MCA causes minimal impact on cerebral tissues, and the ischemic damage is only due to the occlusion of the cerebral artery. Our results provide clear evidence that the surgery required to induce the MCA occlusion induces protein expression changes in the brain, which supports the needed of including sham groups in the experimental designs of preclinical studies. These sham controls are no neutral, and only through appropriate preliminary experiments can be determined how these interventions interfere with the primary purpose of the study.

## MATERIALS AND METHODS

### Animal care and housing

The protocols for rodent assays were approved by the University Clinical Hospital of Santiago de Compostela and the Health Research Institute of Santiago de Compostela (IDIS) Animal Care Committees under procedure numbers: 15011/2022/003 and 15011/2023/001, according to the European Union (EU) rules (86/609/CEE, 2003/65/CE and 2010/63/EU) and within the ARRIVE guidelines. Male Sprague-Dawley rats (7–8 weeks), with a weight of 250–300 g, were used. Animals were housed at an environmental temperature of 23 °C with 40% relative humidity and had a 12-hour light-dark cycle. Rats were watered and fed *ad libitum*. To minimize stress, the animals were acclimated after arrival at the animal facility for at least 1 week. Surgical procedures and magnetic resonance imaging (MRI) were performed under sevoflurane anesthesia (6% induction and 4% maintenance in a mixture of 70% nitrous oxide and 30% oxygen). Rectal temperature was maintained at 37 ± 0.5 °C using a feedback-controlled heating pad (Neos Biotec, Pamplona, Spain). Glucose levels analyzed before surgery showed similar animal ranging from 180 to 220 mg/dL. At the end of the procedures, rats were sacrificed under deep anesthesia (8% sevoflurane).

### Study groups and surgical procedures

In this study, a total of 9 animals were included. These animals were randomized into the following groups: control group, sham group and ischemic group.

Control group: animals exposed to sevoflurane for 120 min (time corresponding to the one used to perform the surgical procedure and to the ischemia occlusion time used in the sham and ischemic group).

Sham group: animals subjected to the same surgical procedure as ischemic animals, without the intraluminal occlusion of the MCA.

Ischemic group: animals induced with transient focal ischemia by the transient intraluminal middle cerebral artery occlusion (tMCAO) model. Transient focal ischemia was induced by intraluminal MCA occlusion as previously described [14, 15] using commercially available sutures with silicone-rubber-coated heads (350 μm in diameter and 1.5 mm long; Doccol, Sharon, MA, USA). In the ischemic groups, cerebral blood flow was monitored with a Periflux 5000 laser Doppler perfusion monitor (Perimed AB, Järfälla, Sweden) by placing the Doppler probe (model 411; Perimed AB) under the temporal muscle at the parietal bone surface, near the sagittal crest. Once the artery occlusion was achieved, as indicated by Doppler signal reduction, each animal was carefully moved from the surgical bench to the MR system for baseline ischemic lesion assessment using MRI apparent diffusion coefficient (ADC) maps (before treatment administration). MR angiography (MRA) was also performed to ensure that the artery remained occluded throughout the MR procedure and to detect possible arterial malformations [16]. After basal MR analysis, animals were returned to the surgical bench and the Doppler probe was repositioned. Reperfusion was performed 75 min after occlusion onset. In line with our previous studies using same ischemic model, the following exclusion criteria were used [14]: (1) <70% reduction in relative cerebral blood flow during arterial occlusion; (2) arterial malformations, as determined by MRA; (3) baseline lesion volume <35% or >45% of the ipsilateral hemisphere, as measured using ADC maps; (4) absence of reperfusion or prolonged reperfusion (>10 minutes until achieving ≥50% of baseline cerebral blood flow) after filament removal. MRI-T2 scans for infarct assessment were determined at 1day after ischemia.

The sham animals were subjected to the same MRI imaging protocol as described for the ischemic group.

### Magnetic resonance imaging and image analysis

MRI studies were conducted on a 9.4 T horizontal bore magnet (Bruker BioSpin, Ettlingen, Germany) with 12-cm wide actively shielded gradient coils (440 mT/m). Radiofrequency transmission was achieved with a birdcage volume resonator and signal was detected using a four-element arrayed surface coil positioned over the head of the animal. The latter was fixed with a teeth bar, earplugs, and adhesive tape. Transmission and reception coils were actively decoupled from each other. Gradient–echo pilot scans were performed at the beginning of each imaging session for accurate positioning of the animal inside the magnet bore. MRI post processing was performed using ImageJ software (https://imagej.nih.gov/ij/). Infarct volumes were determined from T2 relaxation maps by manually selecting areas of hyperintense T2 signal by a researcher blinded to the animal protocols. Infarct size was indicated as the percentage of ischemic damage with respect to the ipsilateral hemispheric volume, corrected for brain edema. For each brain slice, the total areas of both hemispheres and the areas of infarction were calculated. An edema index was measured by quantifying the midline deviation (MD) calculated as the ratio between the volume of the ipsilateral hemisphere and the volume of the contralateral hemisphere. The actual infarct size was adjusted for edema by dividing the area of infarction by the edema index [mm^3^/MD]. Thereafter, the presented infarct volume was calculated as follows: (infarct volume [mm^3^/MD]/ipsilateral hemispheric area [mm^3^]) × 100. These procedures have been used repeatedly in the literature to measure and evaluate stroke outcome in experimental models [15, 17, 18]

#### MR angiography

non-invasive angiography was evaluated with the time-of-flight magnetic resonance angiography (TOF-MRA) as reported previously [14, 16]. TOF-MRA scans were performed with a 3D-Flash sequence with an ET = 2.5 ms, RT =15 ms, FA= 20°, NA = 2, SW = 98 KHz, 1 slice of 14 mm, 30.72 × 30.72 × 14 mm^3^ FOV (with saturation bands to suppress signal outside this FOV), a matrix size of 256 × 256 × 58 (resolution of 120 µm/ pixel × 120 µm/pixel × 241 µm/pixel) and implemented without fat suppression option.

#### MRI T2-maps

ischemic lesions were determined from T2-maps calculated from T2-weighted images acquired 24 hours, 7 and 14 days after the onset of ischemia using a MSME sequence: with an ET = 9 ms, RT = 3 seconds, 16 echoes with 9 ms echo spacing, flip angle (FA) = 180°, NA = 2, spectral bandwidth (SW) = 75 KHz, 14 slices of 1 mm, 19.2 × 19.2 mm^2^ FOV (with saturation bands to suppress signal outside this FOV), a matrix size of 192 × 192 (isotropic in-plane resolution of 100 µm/ pixel x×100 µm/pixel) and implemented without fat suppression option.

### Qualitative and quantitative proteomic analysis in brain tissue

#### Perfusion and Tissue processing

animals were deeply anesthetized with sevoflurane (6% in a mixture of 70% NO_2_ and 30% O_2_) and transcardially perfused with 100 mL of 0.1 M PBS (pH 7.4). Brains were carefully removed from the skull and sectioned at 2 mm thick using a matrix. Tissue was stored at 80 °C for further analysis.

#### Protein extraction and digestion

frozen tissue (100 mg) from the different brain samples was homogenized in 300 μl RIPA buffer [200 mmol/L Tris/HCl (pH 7.4), 130 mmol/L NaCl, 10% (v/v) glycerol, 0.1% (v/v) SDS, 1% (v/v) Triton X-100, 10 mmol/L MgCl2] with antiproteases and anti-phosphatases (Sigma-Aldrich, St. Louis, MO, USA) in a TissueLyser II (Qiagen, Tokyo, Japan). The homogenate was centrifuged at 14,000 *g* at 4 °C for 20 min. Protein concentration was measured using a RC-DC kit (Biorad Lab., Hercules, CA, USA) according to the manufacturing protocol. Protein aliquots of 100 μg were concentrated in an SDS-PAGE single band [19, 20] and submitted to a manual digestion as described elsewhere [20]. Finally, after peptide extraction using 50% (v/v) ACN/0.1% (v/v) TFA (x3) and ACN (x1), peptides were pooled, concentrated in a SpeedVac and stored at −20 °C.

#### Qualitative (LC-MS/MS) Analysis

For protein identification, digested peptides from each sample were separated using reverse phase chromatography. The gradient was developed using a micro liquid chromatography system (Eksigent Technologies nanoLC 400, Sciex, Redwood City, CA, USA) coupled to a high-speed Triple TOF 6600 mass spectrometer (Sciex, Redwood Int. J. Mol. Sci. 2021, 22, 226 18 of 22 City, CA, USA) with a microflow source. The analytical column used was a Chrom XP C18 silica-based reversed-phase column (150 0.30 mm) with a 3mm particle size and 120Å pore size (Eksigent, Sciex Redwood City, CA, USA). The trap column was a YMCTRIART C18 (YMC Technologies Teknokroma Analítica, Barcelona, Spain), with a 3mm particle size and 120Å pore size, that was switched on-line with the analytical column. Data were acquired with a TripleTOF 6600 System (Sciex, Redwood City, CA, USA) using a data-dependent workflow (DDA). The micro-pump generated a flow-rate of 5 µl/min and was operated under gradient elution conditions, using 0.1% formic acid in water as mobile phase A, and 0.1% formic acid in acetonitrile as mobile phase B. Peptides were separated using a 90 minutes gradient ranging from 2% to 90% mobile phase B.

Data acquisition was performed by a TripleTOF 6600 System (Sciex, Foster City, CA) using a data-dependent analysis (DDA) workflow. Source and interface conditions were the following: ionspray voltage floating (ISVF) 5500 V, curtain gas (CUR) 25, collision energy (CE), 10 and ion source gas 1 (GS1) 25. Instrument was operated with Analyst TF 1.7.1 software (Sciex, USA). Switching criteria was set to ions greater than mass to charge ratio (m/z) 350 and smaller than m/z 1400 with charge state of 2–5, mass tolerance of 250 ppm and an abundance threshold of more than 200 counts per second (cps). Previous target precursor ions were excluded for 15 s. The instrument was automatically calibrated every 4 hours using tryptic peptides from PepCalMix as external calibrant.

### Data analysis

After MS/MS analysis (MS2 data), data files were processed using ProteinPilot^TM^ 5.0.1 software from Sciex, which uses the algorithm ParagonTM for database search and ProgroupTM for data grouping. Data was searched using rattus specific Uniprot database, specifying iodoacetamide at cysteine alkylation as variable modification and methionine oxidation as fixed modification. False discovery rate was performed using a non-lineal fitting method, displaying only those results that reported a 1% Global false discovery rate or better[21].

#### Generation of the reference spectral library

a pool of each group was analyzed by a shotgun data-dependent acquisition (DDA) approach. The samples were separated in a micro-LC system Ekspert nLC425 (Eksigen, Dublin, CA, USA) using an Chrom XP C18 150 mm × 0.30 mm, 3 mm particle size and 120 Å pore size (Eksigen, Dublin, CA, USA) at a flow rate of 10 μL/min, using as solvent A water, 0.1% formic acid (FA) and solvent B acetonitrile (ACN), 0.1% FA. The peptide separation gradient was from 5% to 95% B for 30 min, 5 min at 90% B and, finally, another 5 min at 5% B for column equilibration, for a total time of 40 min. Coupled with the LC was a hybrid quadrupole-TOF mass spectrometer, 6600+ (SCIEX, Framingham, MA, USA). Using the mass spectrometer, a 250 ms survey scan was performed from 400 to 1250 m/z followed by MS/MS experiments from 100 to 1500 m/z (25 ms of acquisition time) for a total cycle time of 2.8 s. The fragmented precursors were added to the dynamic exclusion list for 15 s, any ion with charge +1 was excluded from the MS/MS analysis. The protein identification was performed using ProteinPilot software v.5.0.1. (SCIEX, Framingham, MA, USA) using a Rattus Norvegicus or Human (to detect human rGOT administered to the rats) specific Uniprot Swiss-Prot database. The false discovery rate (FDR) was set to 1 for peptides and proteins with a confidence score above 99% [21].

#### Quantification by SWATH and data analysis

the quantitative proteomic analysis was made by SWATH method in a hybrid quadrupole-TOF mass spectrometer, 6600+ (SCIEX, Framingham, MA, USA) (Sciex) as was previously described by our group [22–24]. The SWATH–MS acquisition was performed using an IDA (independent data analysis) method. Four micrograms of protein from each individual sample were subjected to chromatographic separation, as described before. The SWATH method is based on repeating a cycle that consisted of the acquisition of 100 TOF MS/MS scans (400 to 1500 m/z, high sensitivity mode, 50 ms acquisition time) of overlapping sequential precursor isolation windows of variable width (1 m/z overlap) covering the 400 to 1250 m/z mass range with a previous TOF MS scan (400 to 1500 m/z, 50 ms acquisition time) for each cycle. Total cycle time was 6.3 s. For each sample set, the width of the 100 variable windows was optimized according to the ion density found in the DDA runs using a SWATH variable window calculator worksheet from Sciex.

The targeted data extraction of the fragment ion chromatogram traces from the SWATH runs was performed by PeakView (version 2.2) using the SWATH Acquisition MicroApp (version 2.0). This application processed the data using the spectral library created from the DDA analysis data loading over this library the individual samples acquired using a SWATH method. In order to obtain the peak areas, up to ten peptides per protein and seven fragments per peptide were selected based on signal intensity; any shared and modified peptides were excluded from the processing.

The integrated peak areas (SWATH areas) were directly exported to the MarkerView software (AB SCIEX) for relative quantitative analysis. MarkerView uses processing algorithms that accurately find chromatographic and spectral peaks direct from the raw SWATH data. First the integrated peak areas were normalized using MLR normalization or Suma total areas, depending on the analysis performed; then, unsupervised multivariate statistical analysis, using principal component analysis (PCA), was performed to compare the data across the samples using scaling. A Student’s t-test analysis using the MarkerView software was performed for comparison among the samples. The deregulated proteins were selected using p-value < 0.05 and FC >1.5 or < 0.8 as cut-off. The individual values of SWATH areas per protein and sample were used to perform the blox pots.

#### Protein functional enrichment and network analysis

the differentially regulated proteins will be subjected to functional analysis and interpreted through various open access bioinformatics tools for analyzing biological information related to molecular functions, biological processes, cellular components, protein classes, pathways, and networks among the large and complex datasets. For functional enrichment and interaction network analysis we will use FunRich (http://funrich.org/index.html). FunRich uses hypergeometric tests, BH and Bonferroni [25].

